# Molecular structure and function of myelin protein P0 in membrane stacking

**DOI:** 10.1101/395491

**Authors:** Arne Raasakka, Salla Ruskamo, Julia Kowal, Huijong Han, Anne Baumann, Matti Myllykoski, Anna Fasano, Rocco Rossano, Paolo Riccio, Jochen Bürck, Anne S. Ulrich, Henning Stahlberg, Petri Kursula

**Affiliations:** Department of Biomedicine, University of Bergen, Bergen, Norway; Faculty of Biochemistry and Molecular Medicine & Biocenter Oulu, University of Oulu, Oulu, Finland; Center for Cellular Imaging and NanoAnalytics (C-CINA), Biozentrum, University of Basel, Basel, Switzerland; Institute of Molecular Biology and Biophysics, Department of Biology, ETH Zurich, Switzerland; Division of Psychiatry, Haukeland University Hospital, Bergen, Norway; Department of Biosciences, Biotechnologies and Biopharmaceutics, University of Bari, Bari, Italy; Department of Sciences, University of Basilicata, Potenza, Italy; Institute of Biological Interfaces (IBG-2), Karlsruhe Institute of Technology, Karlsruhe, Germany; Institute of Organic Chemistry, Karlsruhe Institute of Technology, Karlsruhe, Germany

**Keywords:** Myelin protein zero, intrinsic disorder, integral membrane protein, lipid phase behaviour, peripheral nervous system, compact myelin

## Abstract

Compact myelin forms the basis of nerve insulation essential for higher vertebrates. Dozens of myelin membrane bilayers undergo tight stacking, and in the peripheral nervous system, this is partially enabled by myelin protein zero (P0). Consisting of an immunoglobulin (Ig)-like extracellular domain, a single transmembrane helix, and a cytoplasmic extension (P0ct), P0 harbours an important task in ensuring the integrity of compact myelin in the extracellular compartment, referred to as the intraperiod line. Several disease mutations resulting in peripheral neuropathies have been identified for P0, reflecting its physiological importance, but the arrangement of P0 within the myelin ultrastructure remains obscure. We performed a biophysical characterization of recombinant P0ct. P0ct contributes to the binding affinity between apposed cytoplasmic myelin membrane leaflets, which not only results in fluidity changes of the bilayers themselves, but also potentially involves the rearrangement of the Ig-like domains in a manner that stabilizes the intraperiod line. Transmission electron cryomicroscopy of native full-length P0 showed that P0 stacks lipid membranes by forming antiparallel dimers between the extracellular Ig-like domains. The zipper-like arrangement of the P0 extracellular domains between two membranes explains the double structure of the myelin intraperiod line. Our results contribute to the understanding of PNS myelin, the role of P0 therein, and the underlying molecular foundation of compact myelin stability in health and disease.

## Introduction

Myelin enwraps axonal segments in the vertebrate nervous system, accelerating nerve impulse propagation as well as providing trophic and mechanical support to fragile neuronal processes^1^. The insulative nature of myelin arises from its water-deficient structure, compact myelin, where plasma membranes are stacked upon each other and adhered together by specific proteins^2^. This array of proteins partially differs between the central and peripheral nervous systems (CNS and PNS, respectively), and the disruption of PNS compact myelin has a severe effect on action potential velocity^3^. This manifests as a set of medical conditions, including the peripheral neuropathies Charcot-Marie-Tooth disease (CMT) and Dejerine-Sottas syndrome (DSS). Such diseases are incurable and difficult to treat, and they show significant genetic background, resulting from mutations in proteins that affect the formation or stability of myelin, either directly or indirectly^4-7^. The development of eventual CMT/DSS-targeting remedies is hindered by the lack of basic molecular structural knowledge on the formation and eventual disruption of PNS myelin^8^.

Myelin protein zero (P0; also known as MPZ) is the most abundant protein in PNS myelin^9^. It resides in compact myelin, and spans the myelin membrane *via* a single transmembrane helix with an N-terminal immunoglobulin (Ig)-like domain on the extracellular side of the membrane. A short cytoplasmic tail (P0ct) follows the transmembrane domain^3^. Point mutations in P0 account for 10 – 12% of all dominant demyelinating CMT type 1 cases^10^.

The extracellular Ig-like domain of P0 is a major contributor to the formation of the myelin intraperiod line^11^. Crystal structures of the Ig-like domain have provided clues into details of membrane adhesion, and one theory involves oligomerization of Ig-like domains from two apposing membranes^12, 13^. This would explain the roughly 5-nm spacing of the intraperiod line in compact myelin, compared to the 3-nm cytoplasmic compartment, or the major dense line, between two cytoplasmic membrane leaflets^14-16^. Dozens of CMT- and DSS-linked mutations have been reported for the Ig-like domain, signifying its importance in myelination^17^.

At physiological pH, P0ct is a positively charged segment of 69 amino acids, predicted to be disordered in solution^3^. The central part of P0ct (amino acids 180-199 of mature human P0 isoform 1) contains an immunodominant, neuritogenic segment, which can be used to generate animal models with experimental autoimmune neuritis (EAN)^18^. It is noteworthy that most CMT-linked point mutations in P0ct are localized in this region^17, 19-22^. P0ct interacts with lipid membranes, and it gains a significant amount of secondary structure upon binding^23-25^. P0ct aggregates negatively charged lipid vesicles ^23^, suggesting that P0ct might harbour a similar membrane-stacking function as myelin basic protein (MBP)^16^ and peripheral myelin protein 2 (P2)^26^. However, the tertiary conformation of P0ct and details of its lipid binding are not fully understood, and the potential function of P0ct in membrane stacking remains to be further elucidated.

We characterized human P0ct using several complementary methods, to gain a structural insight into its membrane binding, insertion, and contribution to myelin membrane stacking. Using electron cryomicroscopy (cryo-EM), we observed a zipper-like assembly of bovine full-length P0 in reconstituted membranes. Additionally, we investigated a synthetic P0ct-derived peptide, corresponding to the neuritogenic sequence, (P0ct_pept_) under membrane-mimicking conditions using synchrotron radiation circular dichroism spectroscopy (SRCD) and computational predictions. Our results show that P0ct is likely to be involved in maintaining the stability of compact PNS myelin together with other cytosolic PNS proteins. P0ct may have a role in assembling into an ordered lateral structure deep within the membrane, potentially stabilizing the ultrastructure formed by the Ig-like domains in the extracellular compartment.

## Results

To study the putative function of P0ct as a classical membrane stacking protein, we performed a biophysical characterization, as well as binding experiments with lipids, similarly to MBP ^16^. Additionally, we investigated the stacking behaviour of full-length P0 using cryo-EM, and the folding and orientation of P0ct_pept_ in membranes with SRCD.

### Characterization of P0ct

We purified P0ct to homogeneity using a two-step purification process. P0ct appeared as a single band on SDS-PAGE, presenting a monodisperse profile in size exclusion chromatography (SEC) with an apparent monomeric mass in solution, as shown with SEC-coupled multi-angle light scattering (SEC-MALS), despite its unusual migration in SDS-PAGE (Supplementary Fig. S1a). The obtained mass (7.5 kDa) was very close to the one determined using mass spectrometry (7991 Da), which in turn matched the expected mass based on the primary structure (7990.25 Da).

The folding of P0ct was studied in the presence of different additives using SRCD. The results were as expected based on earlier investigations^24^: P0ct was mostly disordered in solution and gained significant secondary structure content, when introduced to increasing concentrations of 2,2,2,-trifluoroethanol (TFE), as well as n-dodecylphosphocholine (DPC) or sodium dodecyl sulphate (SDS) micelles, but not with n-octylglucoside (OG) or lauryldimethylamine *N*-oxide (LDAO) (Supplementary Fig. S1b).

To investigate the solution behaviour of P0ct, we used small-angle X-ray scattering (SAXS) to reveal that P0ct is monomeric and extended (Fig. 1a-b, Supplementary Fig. S1c, Supplementary Table S1), which was best explained by a dynamic ensemble of elongated conformers, as observed for MBP (Fig. 1c)^16^. Similarly to MBP, P0ct appeared to favour a slightly compacted conformation over a completely extended one.

**Fig. 1.**
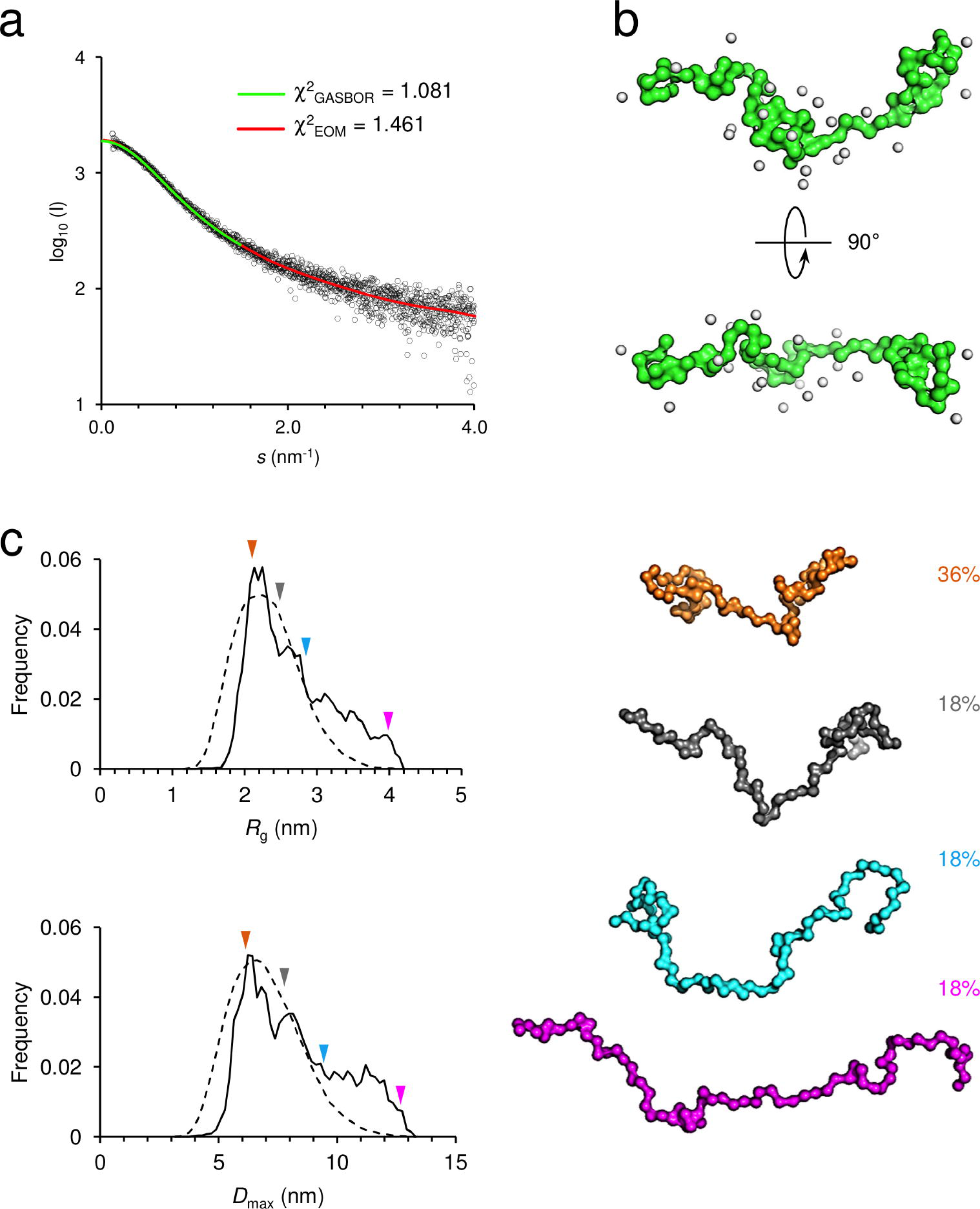
SAXS analysis of P0ct. (a) SAXS data reveals the elongated nature of P0ct. GASBOR and EOM fits are plotted over the entire measurement data with their respective χ^2^ values indicated. (b) The GASBOR *ab initio* model is clearly elongated, measuring up to 8 nm in length. (c) An ensemble of dynamic conformers in solution, presenting wide *R*_g_ and *D*_max_ populations (left), satisfy the SAXS data well. Most abundant conformers by mass are presented as models (right; colored arrows denote the *R*_g_ and *D*_max_ values within the analyzed distributions from EOM).

### P0ct binds irreversibly to lipid structures and influences their properties

A gain in helical structure of P0ct was observed earlier in membrane-mimicking conditions, whereas at increasing lipid concentrations, the formation of β-strands was induced in the presence of high amounts of cholesterol^24^. This could be a physiologically relevant state of folding, which prompted us to investigate the lipid interaction propensity of P0ct in the presence of various lipid compositions by SRCD (Fig. 2a-c). Magnitude differences in the CD signals were detected between different samples, which most likely arose from lipid batch heterogeneity and the resulting differences in protein-lipid turbidity, resulting in light scattering and lowered signal intensity. This did not compromise data interpretation, as the relevant data for each comparison were acquired at the same time. Therefore, the SRCD traces in Fig. 2a-c should be compared only within panels. We observed folding of P0ct in the presence of negatively charged small unilamellar vesicles (SUVs), whereas neutral SUVs displayed no gain in secondary structure content (Fig. 2a). Fully saturated lipid tails, in this case a 1:1 molar mixture of dimyristoylphosphatidylcholine (DMPC) and dimyristoylphosphatidylglycerol (DMPG), enhanced the folding notably, indicating a specific effect by the lipid tails. To check whether other lipid species could influence the folding, we included 10% (w/w) cholesterol, dimyristoylphosphatidylethanolamine (DMPE), or sphingomyelin (SM) in DMPC:DMPG (1:1) (Fig. 2b). Cholesterol and SM appeared to slightly favour folding, suggesting that P0ct is sensitive to fluidity differences within the membrane, as reported earlier^24^. Additionally, these interactions seem specific from a molecular point of view, as changing the fluidity phase of DMPC:DMPG (1:1) *via* temperature scanning did not influence the folding of P0ct (Fig. 2c). SM and cholesterol are known to interact^27^, and together, they contribute to the formation of lipid rafts, in which major PNS myelin proteins localize^28^. These include P0, as well as its known interaction partner peripheral myelin protein 22 (PMP22)^29, 30^, for which the presence of SM is absolutely required for successful reconstitution into functional model systems^31^.

**Fig. 2.**
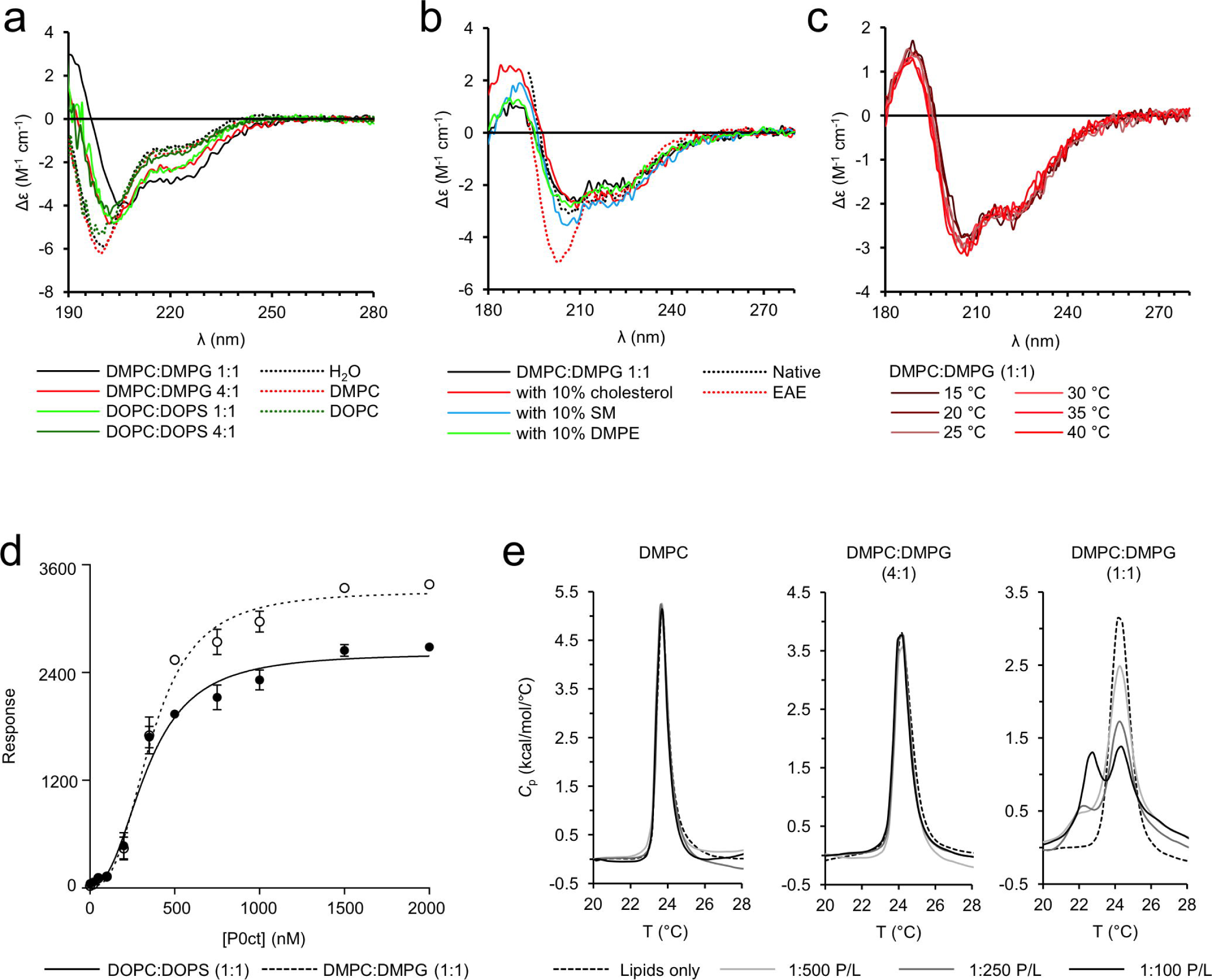
Folding and lipid binding properties of P0ct. (a) While P0ct remains disordered in the presence of neutral lipid bodies, a large gain in regular secondary structure content is evident especially in the presence of DMPC:DMPG (1:1). (b) P0ct appears to fold slightly differently in the presence of vesicles containing cholesterol or SM when mixed with a DMPC:DMPG (1:1) composition. Native and EAE-like lipid compositions (compositions based on previously published data^32^) present a different degree of folding. (c) P0ct retains its folding in SRCD around the typical DMPC:DMPG (1:1) lipid phase transition temperature. (d) P0ct binds immobilized lipid vesicles in SPR. Error bars represent standard deviation. (e) The lipid tails of LMVs with a strong negative surface net charge exhibit a second endothermic transition at 22.5 °C upon the addition of P0ct.

In our model lipid compositions, we did not observe structures rich in β-sheets, and to study more physiologically relevant conditions, we measured SRCD spectra of P0ct in the presence of a native myelin-like cytoplasmic leaflet composition DMPC:dimyristoylphosphatidylserine (DMPS):DMPE:SM:cholesterol (25.9:7:29:6.2:31.6), as well as an experimental autoimmune encephalomyelitis (EAE)-like composition DMPC:DMPS:DMPE:SM:cholesterol (20.1:7.4:32.9:2.2:37.4) (Fig. 2b)^32^. P0ct gained a high amount of secondary structure in the presence of native myelin-like lipids, but did not display enriched amounts of β-structures. The folding was clearly diminished in EAE-like lipids, and a key influencing factor might be the different phase behaviour of the two lipid compositions, which has been described before^33^. Considering the composition, the largest relative difference is the much lower fraction of SM in the EAE-like lipids. It seems unlikely that P0ct would adopt a conformation composed of predominantly β-sheet in myelin. Additionally, judging from the intensity of the observed double minima at 209 and 222 nm, the helical folding of P0ct in our model lipids matches the native-like lipid composition better than the EAE composition, validating our simplified model system for further studies.

The folding of P0ct in the presence of lipid vesicles is a strong indicator of binding, which prompted us to study the binding kinetics and affinity using surface plasmon resonance (SPR; Fig. 2d). P0ct irreversibly associated with immobilized large unilamellar lipid vesicles (LUVs), and comparison of saturated and unsaturated lipid tails revealed only differences in maximum response, reflecting the amount of bound P0ct, while the affinity (*K*_d_) remained similar (Table 1). These results suggest that saturated lipids can incorporate more P0ct; furthermore, P0ct gained more secondary structure in the presence of saturated lipids (Fig. 2a). We were interested in the underlying kinetics of association, but similarly to MBP, we observed a complex association pattern^16^, which could not be analyzed confidently using simple binding models (Supplementary Fig. S2, Supplementary Table S2).

**Table 1.** SPR fitting parameters.

| **Lipid composition** | **$R_{hi}$** | **$R_{lo}$** | **$A_1$** | **$A_2$** | **$R^2$** |
| --- | --- | --- | --- | --- | --- |
| DOPC:DOPS (1:1) | $2617 \pm 80.9$ | $29.1 \pm 64.7$ | $337.2 \pm 20.7$ | $2.44 \pm 0.31$ | 0.9800 |
| DMPC:DMPG (1:1) | $3309 \pm 71.4$ | $19.8 \pm 58.4$ | $363.0 \pm 15.0$ | $2.76 \pm 0.26$ | 0.9893 |

Having observed a similar lipid tail-dependent binding behaviour for P0ct as for MBP, we employed differential scanning calorimetry (DSC) to check for any thermodynamic changes in lipid tail behaviour of large multilamellar vesicles (LMVs) in the presence of P0ct (Fig. 2e). While the phase behaviour of neutral lipids remained unaffected by the presence of P0ct, displaying only the ∼24 °C peak typical for dimyristoyl compounds^34^, the inclusion of high amounts of negatively charged DMPG allowed us to observe an additional endothermic phase transition peak around 22.5 °C. The newly stabilized lipid population at 22.5 °C increased in intensity with the addition of more P0ct, indicating direct binding to vesicles and concentration dependency. Note that P0ct did not display folding differences at temperatures around the observed phase transitions (Fig. 2c), suggesting that P0ct does not seem to exist in several temperature-dependent conformational states. P0ct seems to directly interact with lipid tails and act as a membrane fluidity modulator, by lowering the bilayer phase transition temperature.

### P0ct forms an ordered system within a lipid membrane

Since we confirmed the binding of P0ct to lipid structures, the next step was to investigate its putative membrane stacking function. We used turbidimetry to indirectly quantify vesicle aggregation by P0ct (Fig. 3a). P0ct induced turbidity in the presence of several lipid mixtures with net negative surface charge.

**Fig. 3.**
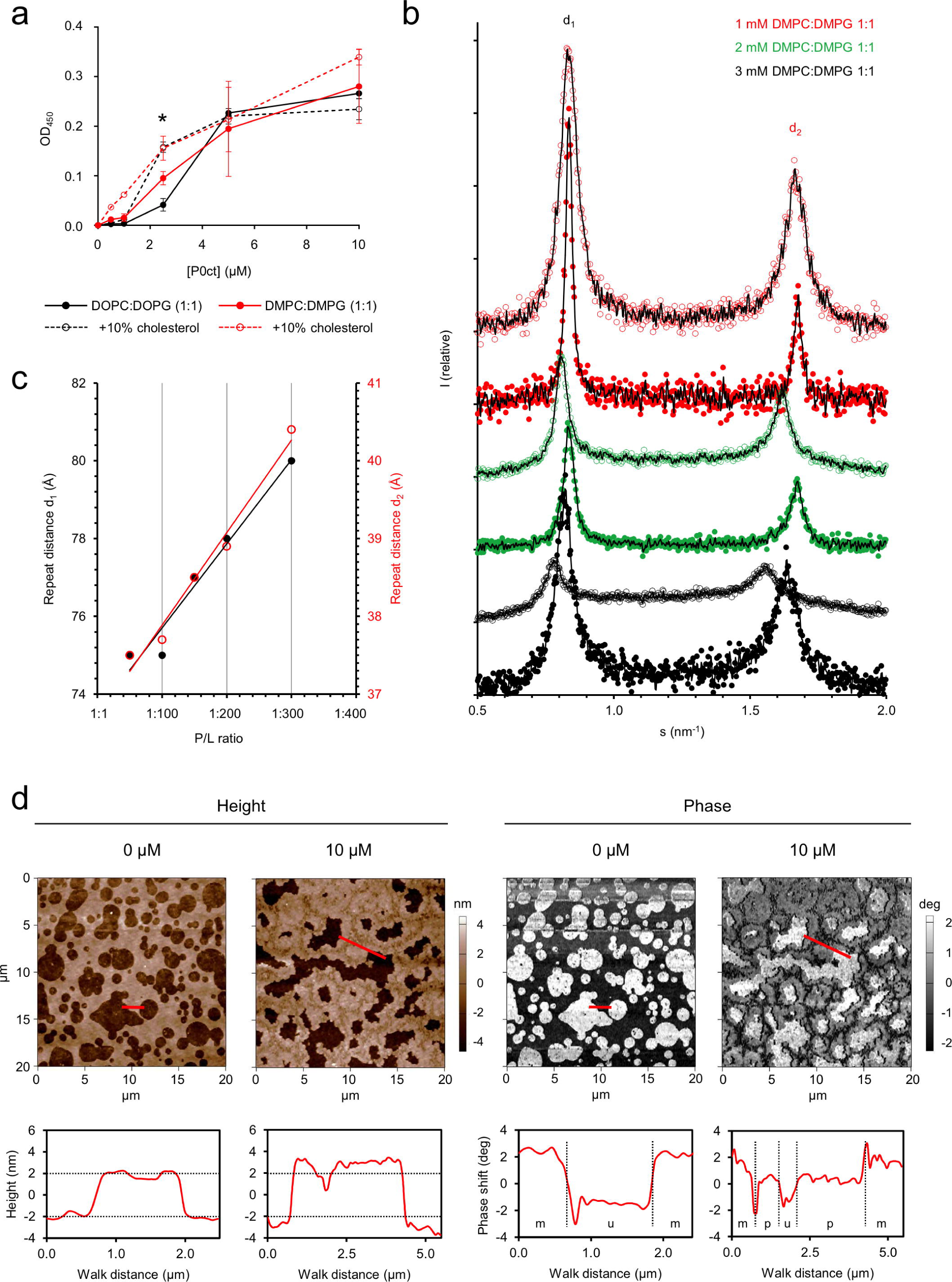
P0ct modulates the properties of lipid structures. (a) The turbidity caused by the presence of P0ct in SUVs somewhat changes depending on the lipid tail saturation degree or the presence of cholesterol. Error bars represent standard deviation. (b) Mixing P0ct with DMPC:DMPG (1:1) SUVs result in X-ray diffraction patterns that display two major Bragg peaks per dataset within the measured *s*-range. Added P0ct concentrations of 10 and 20 µM are shown as open and filled markers, respectively. A moving average (black line) has been plotted over each dataset, all of which have been offset for clarity. None of the data has been scaled in respect to one another. (c) The mean repeat distances, calculated from the *s*-values corresponding to the intensity summit of each Bragg peak, plotted as a function of P/L ratio. Linear fits are shown as solid lines for both d_1_ and d_2_ distributions. (d) In AFM, P0ct accumulates into supported DOPC:DOPS (1:1) bilayers without spontaneous induction of myelin-like membrane stacking, but increases the local thickness of the membrane by as much as 2 nm. The phase difference of the membrane indicates that the membrane rigidity increases in these areas. Height and phase difference graphs are plotted below the images, with the walk path indicated in red. Annotations in phase image walk graphs: m, mica; u, lipid bilayer; p, P0ct-embedded bilayer. Dashed horizontal and vertical dividers have been added to aid comparison.

The turbidimetric experiments strongly indicated that P0ct induced vesicle aggregation. To further elucidate the potential order within the system, we subjected protein-lipid samples to small-angle X-ray diffraction (SAXD) (Fig. 3b-c). While vesicles alone did not show Bragg peaks in the X-ray scattering profile (not shown), P0ct induced two intensive Bragg peaks, which changed their position in momentum transfer (*s*) in a nearly linear fashion as a function of protein-to-lipid (P/L) ratio, indicating concentration dependency in the formation of order and membrane spacing.

While the vesicle-based experiments provided indirect evidence of membrane stacking, we used atomic force microscopy (AFM) to determine whether P0ct can spontaneously stack membranes, similarly to MBP and P2^26^ (Fig. 3d). However, P0ct did not form stacked systems even at relatively high concentrations, rather appearing to accumulate on and increase the thickness and rigidity of the membranes. This difference to turbidimetric and SAXD experiments drove us to exploit EM to check whether aggregated vesicles form in the presence of P0ct (Fig. 4a-d). Surprisingly, the EM data revealed an absence of vesicle aggregation and clear membrane stacks; the difference to control samples was the formation of large vesicular bodies, which likely form through P0ct-mediated fusion, at most tested P/L ratios. At high protein concentrations, we observed 10–20-nm thick filament/tubule-like assemblies, seemingly formed of tightly adhered membranes. This is an indication of ordered protein-membrane structures, although such filamentous structures are not present in endogenous myelin. It appears that high enough P0ct concentrations may affect membrane bilayer curvature, which could be relevant for myelination. It should be noted that in these experiments, the required P/L ratio for P0ct was roughly an order of magnitude higher compared to MBP to form any kind of visibly stacked systems^16^; the concentration of P0 in PNS compact myelin is several-fold higher than that of MBP, on the other hand. The large vesicular structures suggest an unexpected mechanism, which does explain the observed turbidity at moderate P/L ratios; at the same time, the Bragg peaks in SAXD indicate the presence of well-defined repeat distances, possibly originating from local patches of adhered membranes. It is noteworthy that at P/L ratios of 1:50 and 1:100, the Bragg peaks are most intensive, and the repeat distance plateaus at 75.0 Å, suggesting that the most condensed structure has been reached. A repeat distance of 75 Å is in good corroboration with the thickness of cytoplasmic membrane stacks in myelin^15^; this distance is shorter than that observed with MBP and P2 earlier. These results indicate that P0ct can function as a membrane-stacking molecule. However, taking all of our results together, the contribution of P0ct in membrane stacking *per se* is most likely low. Given the high P0ct concentration needed for the formation of large-scale assemblies, its presence could rather enable the formation of a protein lattice within the membrane itself.

**Fig. 4.**
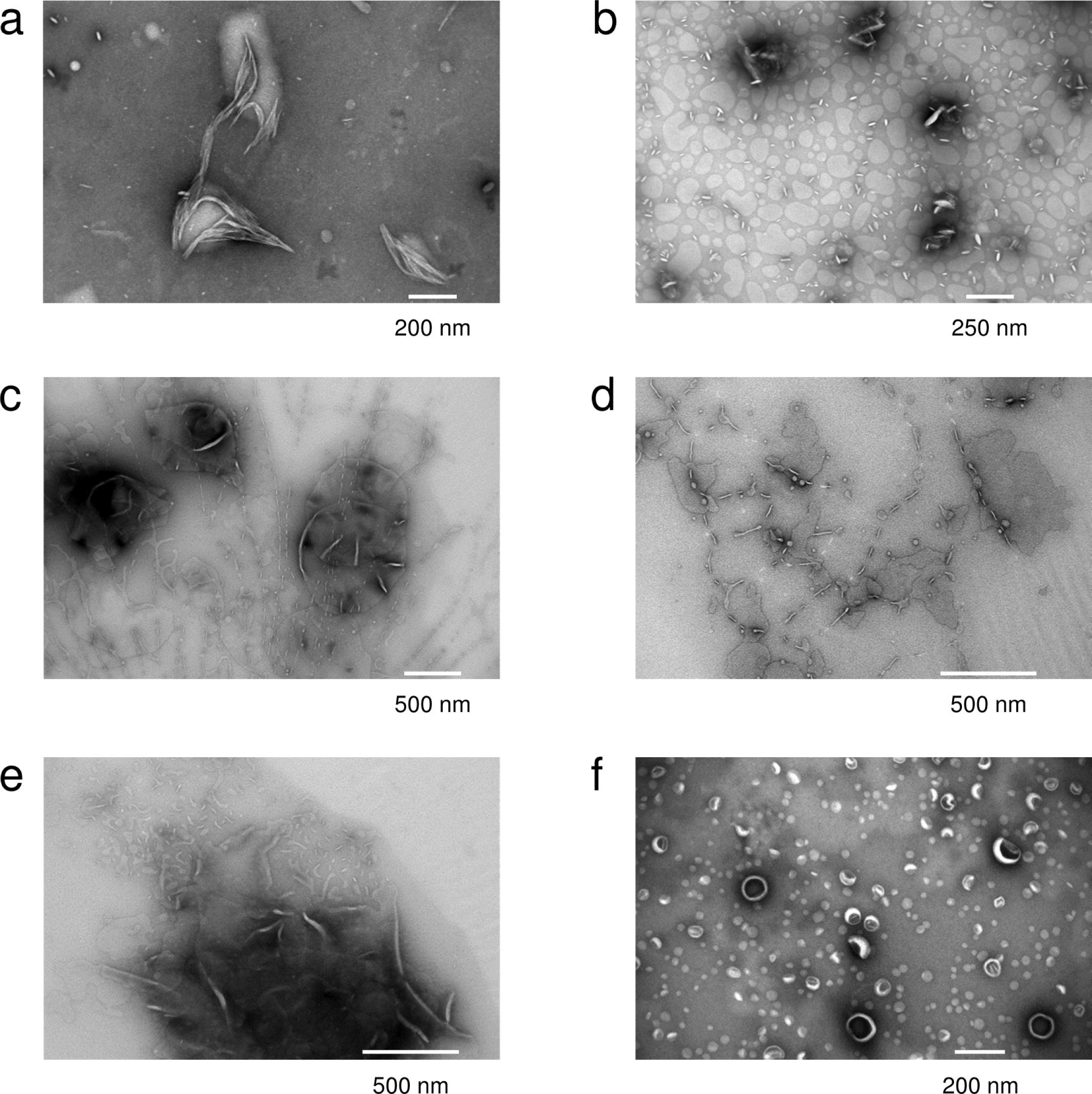
Vesicle fusion and membrane insertion of P0ct. EM concentration series of (a) 1:25, (b) 1:50, (c) 1:100, (d) 1:250 and (e) 1:500 P/L P0ct in the presence of DMPC:DMPG (1:1) vesicles shows the absence of membrane stacks, oppositely to the case of MBP^16^. The effect resembles vesicle fusion and the formation of large lipid bodies. (f) Vesicle control in the absence of P0ct.

### The folding and orientation of P0ct_pept_ in membranes

The P0ct_pept_ peptide has been used to induce EAN in animal models^18^. To shed light on its membrane association and conformation, we studied P0ct_pept_ using SRCD and oriented SRCD (OCD). P0ct_pept_ was disordered in solution and gained secondary structure in the presence of TFE, negatively charged detergent micelles, and somewhat less in negatively charged lipids (Fig. 5a,b). P0ct_pept_ did not fold with neutral lipids, and relatively weakly with negatively charged lipids, except with DMPC:DMPG (1:1). In the latter composition, the gain in secondary structure was notable, and the CD spectrum does not directly resemble classical α-helical or β-sheet lineshape: the observed maximum and minimum at 188 nm and 205 nm, respectively, are close to typical π⟶π* transitions observed in peptides and proteins with mostly α-helical structures. However, the n⟶π* band, normally observed as a minimum at 222 nm in α-helices, is poorly resolved. While the folding of P0ct_pept_ did not gain significant structure in DPC, the negatively charged detergent SDS induced higher secondary structure gain than 70% TFE, in contrast to P0ct and several other proteins^16, 35, 36^, suggesting a role for electrostatic neutralization in the folding of P0ct_pept_.

**Fig. 5.**
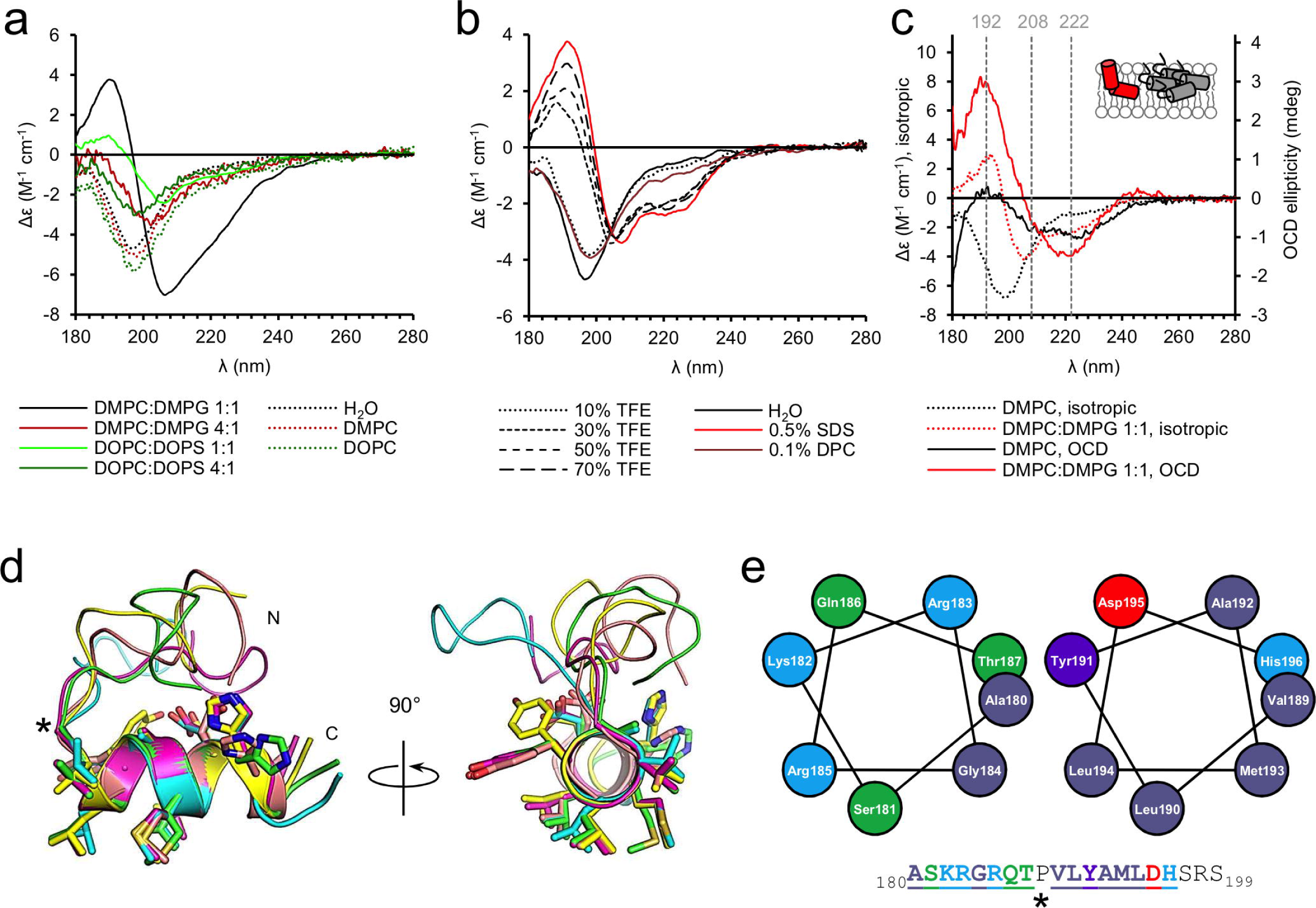
Folding of P0ct_pept_. (a) SRCD data show that P0ct_pept_ only displays folding with negatively charged SUVs. The most notable spectrum is the DMPC:DMPG (1:1) spectrum, which does not resemble classical helical content, nor β-sheets. (b) The effect of TFE and detergents on the conformation of P0ct_pept_. While P0ct_pept_ only displays marginal folding with DPC, SDS induces an even higher secondary structure gain than 70% TFE, indicating the major importance of electrostatics in the folding of P0ct_pept_. (c) OCD data of P0ct_pept_ in DMPC and DMPC:DMPG (1:1). Isotropic spectra are shown for reference. The DMPC OCD data is weak, typical for an aggregated sample, while also showing traces of helical content. The DMPC:DMPG (1:1) data displays clear folding and a more perpendicular orientation within the membrane. A simple cartoon has been drawn for clarity for both OCD datasets. Colors match the shown data. (d) Superposition of five predicted structures of P0ct_pept_ reveal the N-terminus likely to be disordered, followed by a kinking Pro (black asterisk), and a helical C-terminal segment. (e) Helical wheel projections of the basic N-terminal half (left) and the hydrophobic half (right) of P0ct_pept_. The residues are colored based on chemical properties with matched indications in the peptide sequence (down). The underlined segments are shown in the projections. The potentially helix-breaking Pro has been indicated with an asterisk.

To shed light on the orientation of P0ct_pept_ in membranes, we prepared stacked membranes with P0ct_pept_ on planar quartz glass substrates at high relative humidity (>97%). The samples were oriented perpendicular to the incident synchrotron radiation beam, and we collected OCD spectra of P0ct_pept_ in DMPC and DMPC:DMPG (1:1) membranes (Fig. 5c), which allows the investigation of helix orientation within oriented membranes^37, 38^. Comparison of the OCD spectra with isotropic spectra revealed folding in both cases. The DMPC OCD spectrum is weak, but presents helical content, with both minima typical for in-plane helices in bilayers present; compared to the isotropic spectrum, this reflects the fact that in OCD, there is much less free water phase and the peptide is forced to bind to DMPC. In the DMPC:DMPG (1:1) data, the spectral intensity is higher, and the minimum typically observed at 208 nm is weakened, indicating that the helical segment of P0ct_pept_ is in a tilted orientation with respect to the lipid surface, although the shift is not strong enough to represent a completely perpendicular helix.

We generated models of P0ct_pept_ to learn more about its potential conformation. The N-terminal, basic half of P0ct_pept_ is likely flexible, whereas the more hydrophobic segment corresponding to residues 189 – 199 folds into a well-defined helix (Fig. 5d,e). Linking the models to the observed OCD spectra, the peptide most likely adopts a mix of orientations in DMPC, whereas in the presence of negatively charged headgroups, the N-terminal half of P0ct_pept_ can form ionic interactions. This could result in charge neutralization and induced folding into a second helical segment, which would adopt a more perpendicular orientation, explaining the measured OCD spectrum. It is likely that the central Pro residue breaks the helix and allows the two halves to adopt different orientations with respect to one another and the membrane. It is possible that as least the central helical segment of P0ct is ‘anchored’ at the level of the charged headgroups, and as such, is likely to give P0ct a specific orientation in the cytoplasmic leaflet of the myelin membrane.

### P0 forms an intermembrane zipper composed of Ig-like domains

To complement the characterization of P0ct above, we investigated membrane stacking by full-length P0, purified from the raft fraction of bovine peripheral myelin^39^. After purification, the protein was embedded in SDS micelles, and after exchange into n-decyl-β-D-maltopyranoside (DM), the protein was monodisperse based on negative stain EM (Fig. 6a). SRCD investigations of P0 in detergents and reconstituted into net-negatively charged lipids produced spectra typical for high structural content (Fig. 6b), and monodispersity was further analyzed using SEC-MALS (Fig. 6c, Supplementary Fig. S3). The oligomeric status of full-length P0 depended on detergent, and in accordance with earlier studies^40, 41^, both tetrameric and dimeric species were observed. The main species in DPC was tetrameric, while LDAO showed the presence of dimeric P0. It is noteworthy that the folding of P0ct was drastically different in these two detergents (Supplementary Fig. S1b), suggesting the possible involvement of P0ct in the oligomerization of P0.

**Fig. 6.**
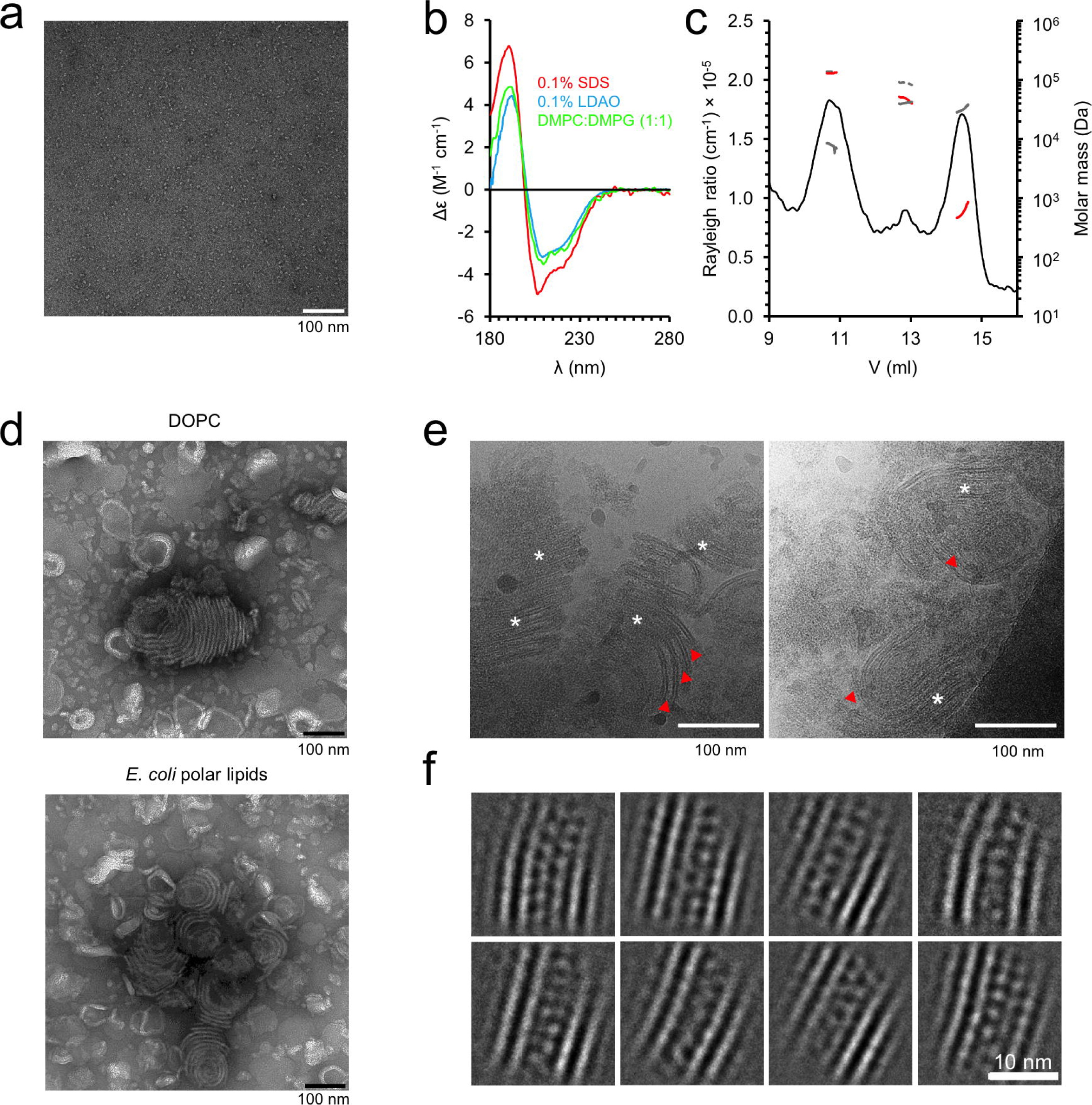
Full-length P0 in a membrane environment. (a) Cryo-EM of full length P0 displays good monodispersity in 0.4% DM. (b) SCRD spectra of full length P0 in detergents and lipids. (c) Analysis of full-length P0 monodispersity and oligomeric state using SEC-MALS. Shown are data measured in DPC. The Rayleigh ratio is shown (black) together with the total mass (gray dash), protein mass (red) and detergent mass for each peak (gray solid). (d) P0 reconstituted into DOPC and *E. coli* polar lipid extract vesicles result in membrane stacks that resemble myelin. (e) Cryo-EM micrographs of *E. coli* polar lipid extract with reconstituted P0. Both curved loosened and planar tight bilayer adhesions are present. In the latter, a significantly higher protein density between the membranes is evident. The loose and tight areas are marked with red arrows and white asterisks, respectively. (f) 2D class averages generated from the tight adhesions of P0 in *E. coli* polar lipids sample cryo-EM micrographs display a zipper of apposing Ig-like domains that interact with each other and settle between the two lipid bilayers.

We reconstituted P0 into SUVs prepared from both DOPC and *E. coli* polar lipids and images the samples with cryo-EM. We observed similar membrane stacking in both lipid compositions, which suggests that the lipid composition plays a minor role in membrane stacking by full-length P0 under these conditions (Fig. 6d-f).

The extracellular domain of P0 was organized between the two apposing lipid bilayers, apparently forming a tightly packed zipper consisting monomers from both apposed membranes (Fig. 6d-f). The thickness of the zipper determined the spatial width, measuring 50 ± 10 Å, between the membrane leaflets – a physiologically relevant value present in natural myelin, as shown by X-ray diffraction of myelin extracts^14, 15^. Each bilayer measured a typical 45-Å thickness. The distance between the apposing P0 monomers was 10-15 Å, if not in direct contact, whereas the spacing between lateral P0 monomers was 25-30 Å. It is realistic to assume that similar distances are present in native myelin, P0 constituting up to 50% of total protein in myelin. The tightly packed, zipper-like arrangement suggests that even if the underlying intermolecular interactions might be relatively weak, the large surface area formed by a P0-enriched membrane can generate a strong force for keeping apposed membranes tightly together.

## Discussion

The cytoplasmic domain of P0 is disordered in solution, and once it irreversibly interacts with negatively charged lipids, it folds and becomes embedded, influencing the phase behaviour of the lipids within the membrane, but also inducing larger-scale events, such as vesicle fusion. P0ct gains helical secondary structure with lipids, rather than the β-sheet-rich structures suggested earlier^24^. This was notable also in a lipid composition that resembles the cytoplasmic leaflet of myelin, and the degree of folding was diminished in an EAE-like lipid composition.

Similarly to MBP^16^, the initial association of P0ct with lipids is governed by electrostatics, suggesting that P0ct possesses similar physicochemical properties as MBP within the myelin environment. However, due to the folding differences observed in SRCD with different lipid tails, correlated with amount of protein bound in SPR, one can speculate that the membrane-bound concentration of P0ct may influence its overall folding population. Already early studies in the 1930s and 1940s have shown that native myelin is an ordered structure diffracting X-rays^42, 43^. P0ct stacks membranes at such a level of order that X-ray diffraction bands are observed, and a repeat distance of 75 Å can be deduced in the multilayers. Lipid-bound MBP, having a high content of PE and low content of bound cholesterol and SM, was shown to similarly generate stacked multilayers with repeat distances between 70-85 Å^44^. Our study provides evidence that P0ct may not harbour a similar role in the formation of the major dense line as MBP, although it is definitely able to bridge charged membranes together at high protein concentrations *in vitro*. In this respect, it is important to remember that *in vivo*, P0ct is an extension of the transmembrane domain of P0, and solidly anchored to the membrane surface, possibly allowing stronger effects on membrane stacking than observed with the P0ct in isolation.

Our cryo-EM experiments on full-length P0 reconstituted into bilayers answer an important question regarding the arrangement of the intraperiod line: the Ig-like domains of P0 organize into apposed dimers, forming a zipper between the two membranes. This is apparently independent of lipid composition, and the interaction is mediated directly through the Ig-like domains. The data clearly indicate that the Ig-like domain of P0 from one membrane does not bind directly to the apposing membrane, but two rows of Ig-like domains are present in the extracellular compartment. This arrangement explains the double structure observed at the intraperiod line in high-resolution electron micrographs^45^. Full-length P0 contains the lipids typical of raft microdomains and N-linked glycan structures belonging to hybrid types^39^. The glycan moieties present in P0 might have a role in the formation of complex structures and the partitioning in the raft microdomains of PNS myelin.

From our data, it is not possible to observe high-resolution molecular details of dimerization. To understand P0 assembly in myelin, the crystal structure of the P0 extracellular domain can be used. As discussed in the original publication^12^, the packing of the P0 Ig-like domain in the crystal state provides a potential side-by-side dimerization scheme with one N-terminal β-strand of each apposing dimer forming hydrogen bonding in an antiparallel fashion (Fig. 7). While the dimer is supposedly fairly unstable, it also reflects in our EM data as loosened areas, where the two monomers appear to be out of contact, mostly in areas displaying increased membrane curvature (Fig. 6e). In this assembly, the C terminus of each Ig-like domain monomer would directly face the myelin membrane. The single 26-residue transmembrane helix of P0 is predicted to be roughly 4.5 nm long, able to span a single lipid bilayer. The Ig-like domain is hovering on the membrane outside surface at the start of the helix.

**Fig. 7.**
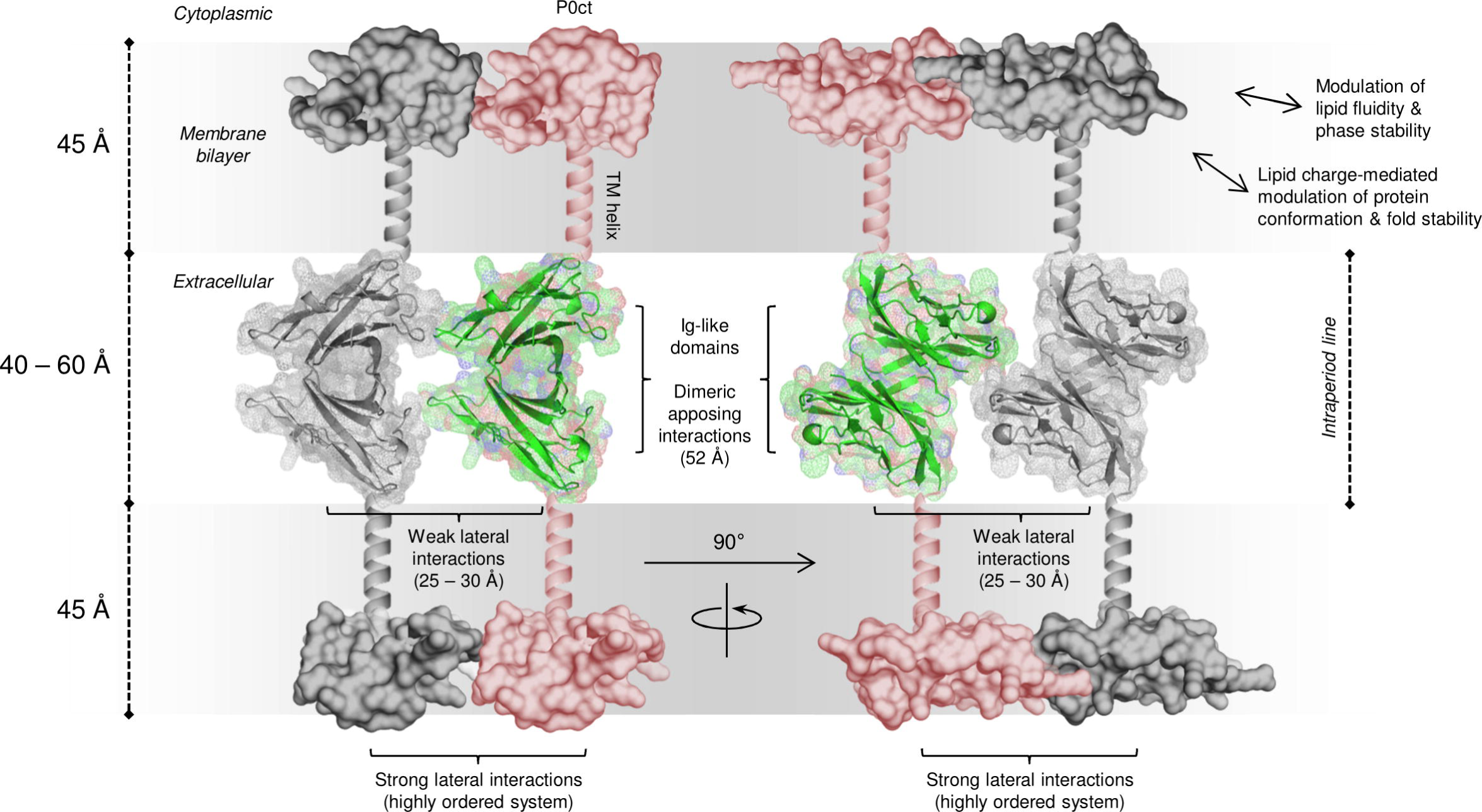
Model for the ultrastructure of the intraperiod line. Potential arrangement of P0 in the intraperiod line and lipid bilayers. Antiparallel apposing Ig-like domains interact weakly with each other through a β-strand stretch (based on crystal contacts in PDB ID 1neu^12^) and form the basis of membrane stacking in the extracellular space. This arrangement results in the C-terminal end of the crystal structure to face the membrane, which logically would turn into a transmembrane helix. Additionally, the size of the antiparallel dimer is similar to the width of the intraperiod line. While the Ig-like domain monomers are not in close contact with one another, P0ct anchors onto the cytoplasmic phospholipid headgroups through electrostatic interactions, gaining folding and potentially forming a stable, ordered lattice within the membrane, which forces P0 molecules to adjacency, including the extracellular domains. Additionally, P0ct changes the properties of the lipid bilayers themselves, offering stiffness and stability to PNS compact myelin. The given lengths in brackets match the measured dimensions from the crystal structure, cryo-EM micrographs, and SAXD.

While P0 dimerizes through the Ig-like domain monomers, the adjacent dimers seem to be separated from one another in the lateral dimension. Therefore, the lateral architecture is likely to be achieved by other factors than the Ig-like domains alone, perhaps through contributions by P0ct on the cytoplasmic face. Our diffraction studies suggest that P0ct is capable of forming higher order structures with membranes, indicating stable molecular interactions (Fig. 7). One could speculate that the function of P0ct is twofold: Firstly, P0ct affects the lateral organization of P0 molecules within the membrane, allowing the formation of organized patches of P0 that together form a strong interaction that keeps the apposing bilayers together. This would affect the orientation of the Ig-like domain to allow the subsequent ‘zipper structure’. Secondly, P0ct influences the fluidity of the membrane itself, potentially by directly associating with certain lipid species, including saturated lipid tails, cholesterol, and SM, which influence membrane phase behaviour that favours productive stacking and the stability of myelin. This could also contribute to the lateral organization of P0 and its interaction partners, such as PMP22^29, 30^. Similarly to P0, mutations in PMP22, many of which are located in transmembrane helices, are hallmark features of CMT and DSS^46^. Changes in the properties and conformation of proteins inside the myelin membrane can have a significant impact on myelin morphology and disease etiology.

Our studies involving the folding of P0ct and P0ct_pept_ suggest a strong influence of lipid electrostatics. While both display folding in the presence of negatively charged lipids, P0ct_pept_ appears to gain a tilted orientation in the membrane upon charge neutralization. This neuritogenic segment of P0ct can directly interact with lipid headgroups and fold into a rigid segment, and as the cytoplasmic side of the lipid bilayer is negatively charged, P0ct could adopt a defined orientation within the membrane. Other positively charged regions of P0ct might function in a similar manner. The adoption of a specific orientation in a membrane would aid in the formation of higher-order structures, which in turn could alter the behaviour, morphology, and rigidity of the membrane *per se* –as exemplified by our EM results at high protein concentrations. In general, membrane curving has been linked to phenomena highly relevant to P0 behaviour observed here at the molecular level; these include the asymmetric insertion of proteins to membrane leaflets, as well as the oligomerization of protein monomers into structures stabilizing a curved membrane shape^47^.

The current study sheds light on the role of P0 at both the major dense line and the intraperiod line. If P0ct stabilizes the arrangement of P0 within a membrane, its importance for the integrity of myelin becomes obvious, and hints of potential molecular disease mechanisms arising from mutations within P0ct, many of which are specifically located within the neuritogenic helical sequence. The impact of these mutations has not yet been studied at the molecular level. Once the molecular basis of the disease mutations in P0ct has been solved, more functional aspects of P0ct will unravel, contributing to a better understanding of PNS compact myelin and its relation to CMT disease etiology.

## Experimental procedures

### Cloning, protein expression, and purification

A synthetic gene encoding for the 69 C-terminal residues (amino acids 151-219) of mature human myelin protein zero isoform 1 (UniProt ID: P25189) was transferred into the Gateway donor vector pDONR221 (Life Technologies). A tobacco etch virus (TEV) protease digestion site (ENLYFQG) was added before the gene, and the required *att*B1 and *att*B2 recombination sites before and after the gene, respectively. This entry clone was used to generate an expression clone in the pHMGWA vector^48^, which encodes for an N-terminal His_6_-tag, followed by a segment encoding for maltose-binding protein (MaBP), and finally followed by the inserted P0ct gene itself (His-MaBP-P0ct). A Cys153Leu mutation was included in the construct for two reasons: this cysteine is known to be palmitoylated *in vivo,* and free cysteines in small, disordered proteins are reactive and can easily result in unwanted intermolecular disulfides^36^. While the introduced Leu residue is not comparable to native palmitoylation, it does manifest as a bulky hydrophobic residue, whilst keeping the protein in a soluble state.

His-MaBP-P0ct was expressed in *Escherichia coli* BL21(DE3) using 0.4 mM IPTG induction for 3 hours in LB medium, at 37 °C. After expression, the cells were collected using centrifugation, broken by ultrasonication in Ni-NTA washing buffer (40 mM HEPES, 400 mM NaCl, 20 mM imidazole, pH 7.5) supplemented with an EDTA-free protease inhibitor cocktail (Roche). Purification was performed using Ni-NTA affinity chromatography using standard procedures. Elution was performed using 32 mM HEPES, 320 mM NaCl, 500 mM imidazole, pH 7.5. The eluted protein was pre-dialyzed at 4 °C with constant stirring against 40 mM Tris-HCl, 400 mM NaCl, 1 mM DTT, pH 8.5, before addition of recombinant TEV protease for His-MaBP tag removal^49^. Quantitative proteolysis was achieved by overnight dialysis, which resulted in cleaved P0ct with an additional N-terminal Gly residue. After this, sequential SEC using a HiLoad Superdex 75 pg 16/60 column (GE Healthcare) was used to separate the cleaved protein from any contaminants, TEV protease, and the cleaved tag, resulting in pure, monodisperse P0ct. Depending on downstream application, either 20 mM HEPES, 300 mM NaCl, 1% (w/v) glycerol, pH 7.5 (SEC buffer) or 20 mM HEPES, 150 mM NaCl, pH 7.5 (HBS) was used as size-exclusion buffer.

Full length P0 was obtained from the raft fraction of bovine nerves as described ^39^. The SDS extract was lyophilized until use. The protein was redissolved in water to regain the original buffer conditions and reconstituted into lipid membranes or other detergents through extensive dialysis. For EM imaging, control samples were taken from redissolved, non-reconsituted samples to control for the presence of putative membranes and multilayers from the original tissue source.

### Mass spectrometry

The identity and accurate molecular weight of P0ct were verified by mass spectrometry. In short, the undigested mass of P0ct was determined using ultra-performance liquid chromatography (UPLC) coupled electrospray ionization (ESI) time-of-flight mass spectrometry in positive ion mode using a Waters Acquity UPLC-coupled Synapt G2 mass analyzer with a Z-Spray ESI source.

### Multi-angle light scattering

SEC-MALS was used to determine the monodispersity and molecular weight of P0ct in solution. Chromatography was performed using an Äkta Purifier (GE Healthcare) and a Superdex 75 pg 10/300GL (GE Healthcare) column with 20 mM HEPES, 300 mM NaCl, pH 7.5 as mobile phase. A 250-µg P0ct sample was injected into the column at an isocratic flow of 0.4 ml/min, and light scattering recorded using a Wyatt miniDAWN TREOS instrument. The UV signal recorded at 280 nm was used as concentration source using the extinction coefficient of P0ct (Abs 0.1% = 1.061) calculated using ProtParam^50^. Data were analyzed using the ASTRA software (Wyatt).

Full length P0 was similarly run on SEC-MALS in the presence of different detergent micelles. The running buffer contained 250 mM sodium phosphate (pH 8.0), 1 mM EDTA, and 0.1% of either DPC or LDAO. A Superdex 200 column was used in the experiment. The protein conjugate analysis routine in the ASTRA software was used to obtain the molecular weight of P0 and the detergent in the separated peaks.

### Small-angle X-ray scattering

SAXS data for P0ct were collected from samples at 1.1 – 4.2 mg ml^−1^ in SEC buffer on the EMBL P12 beamline, DESY (Hamburg, Germany). Monomeric bovine serum albumin was used as a molecular weight standard. See Supplementary Table 1 for further details. Data were processed and analyzed using the ATSAS package^51^. GNOM was used to calculate distance distribution functions^52^, and *ab initio* modelling was performed using GASBOR^53^. Ensemble optimization analysis was performed using EOM^54^.

### Vesicle preparation

Cholesterol, DMPC, DMPG, DMPS, DOPC, DOPG and SM were purchased from Larodan Fine Chemicals AB (Malmö, Sweden). DMPE, DOPS and the deuterated d_54_-DMPC and d_54_-DMPG were purchased from Avanti Polar Lipids (Alabaster, Alabama, USA).

Lipid stocks were prepared by dissolving the dry lipids in chloroform or chloroform:methanol (1:1 v/v) at 5-10 mg ml^−1^. All mixtures were prepared from the stocks at desired molar or mass ratios, followed by solvent evaporation under an N_2_ stream and freeze-drying for at least 4 h at −52 °C under vacuum. The dried lipids were either stored air-tight at −20 °C or used directly to prepare liposomes.

Liposomes were prepared by agitating dried lipids with either water or HBS at a concentration of 2-10 mg ml^−1^, followed by inverting at ambient temperature, to ensure that no unsuspended lipids remained in the vessel. MLVs were subjected to freeze-thawing using liquid N_2_ and a warm water bath, with vigorous vortexing afterwards. Such a cycle was performed 7 times. LUVs were prepared by passing fresh MLVs through a 0.1-µm membrane 11 times on a 40 °C heat block. SUVs were prepared by sonicating fresh MLVs. Either probe tip sonicators (a Branson Model 450 and a Sonics & Materials Inc. Vibra-Cell VC-130) or a water bath sonicator with temperature control (UTR200, Hielscher, Germany) were used to clarify the liposome suspensions, while avoiding overheating. All lipid preparations were immediately used for the designated experiments.

### Synchrotron radiation circular dichroism spectroscopy

The synthetic P0ct_pept_ peptide (NH_2_-ASKRGRQTPVLYAMLDHSRS-COOH) was ordered from GenScript as 5 mg lyophilized aliquots, which were dissolved directly in water. Folding predictions of P0ct_pept_ were generated using PEP-FOLD^55^.

Full-length P0 was studied with SDS, LDAO, and DOPC:DOPS. For preparing P0 in detergents, the sample from purification was dialyzed against 10 mM sodium phosphate buffer (pH 8.0) containing either 0.1% SDS or 0.1% LDAO. For reconstitution into lipids, DOPC:DOPS (1:1) were added to 1 mg/ml, and dialysis was carried out against 10 mM sodium phosphate buffer (pH 8.0).

Isotropic SRCD data were collected from 0.2 – 0.5 mg ml^−1^ protein and peptide samples in water on the UV-CD12 beamline at KARA (KIT, Karlsruhe, Germany)^38^ and the AU-CD beamline at ASTRID2 (ISA, Aarhus, Denmark). We used unbuffered conditions to exclude any unwanted effects, like the electrostatic binding of inorganic phosphate to P0ct, which could interfere with protein binding to detergents and phospholipids. Samples containing lipids were prepared right before measurement by mixing P0ct (P/L ratio 1:200) or P0ct_pept_ (P/L ratio 1:25) with freshly sonicated SUVs, followed by degassing in a water bath sonicator at ambient temperature for 5 – 10 min. 100-µm pathlength closed cylindrical cells (Suprasil, Hellma Analytics) were used for the measurements. Spectra were recorded from 170 to 280 nm at 30 °C and truncated based on detector voltage levels as well as noise. After baseline subtraction, CD units were converted to Δε (M^−1^ cm^−1^), using P0 and P0ct concentration determined from absorbance at 280 nm, or by calculating from the stock concentration of the peptide. SDS and TFE were purchased from Sigma-Aldrich and the detergents LDAO, OG, DM, and DPC from Affymetrix. P0ct and P0ct_pept_ were measured several times in the absence of additives, as well as with DOPC:DOPS (1:1) and DMPC:DMPG (1:1) and the observed spectra were deemed reproducible – the observed trends were always the same and the spectral maxima and minima were accurately present at their typically observed wavelengths.

OCD spectra were measured at the UV-CD12 beamline at KARA (KIT, Karlsruhe, Germany)^38^. 10 µg of P0ct was mixed with 200 µg DMPC or DMPC:DMPG (1:1) SUVs to yield 1:50 P/L samples in water, which were carefully dispensed on quartz glass plates (Suprasil QS, Hellma Optik GmbH, Jena, Germany) within a circular (Ø 1.2 cm) area and allowed to dry at ambient temperature. A background sample without peptide was also prepared. The samples were assembled into humidifier chambers containing a saturated K_2_SO_4_ solution, and allowed to swell for 16 hours at 30 °C and over 97% relative humidity. After the swelling-induced formation of oriented lipid bilayers, the sample chambers were mounted on the beamline (swelled sample perpendicular to the incident beam) and allowed to equilibrate to the original 30 °C temperature and >97% relative humidity. Single-scan spectra were recorded from 170 to 280 nm in 0.5 nm steps at sample rotation angles 0, 45, 90, 135, 180, 225, 270, 315°. The eight spectra were averaged and the lipid background spectrum was subtracted from the sample spectrum.

### Differential scanning calorimetry

P0ct was mixed with MLVs in HBS at several protein-to-lipid ratios (1:250 – 1:1000), always containing 160 µM of either DMPC, DMPC:DMPG (4:1), or DMPC:DMPG (1:1), in final volumes of 700 µl. Lipid samples without P0ct were prepared as controls. The samples were incubated at 37 °C for 10 min to ensure thorough protein association with the vesicles, and degassed for 10 min in vacuum with stirring at 10 °C before measurements.

DSC was performed using a MicroCal VP-DSC calorimeter with a cell volume of 527.4 µl. The reference cell was filled with HBS. Each sample was scanned from 10 to 50 °C and again back to 10 °C in 1 °C min^−1^ increments. Baselines were subtracted from sample curves and zeroed between 15 and 20 °C to enable easier cross-comparison. All samples were prepared and measured twice, with the underlying trends being reproducible.

### Surface plasmon resonance

SPR was performed on a Biacore T200 system (GE Healthcare). According to the manufacturer’s instructions, 100-nm LUVs of 1 mM DMPC:DMPG (1:1) and 1 mM DOPC:DOPS (1:1) were immobilized on separate channels on an L1 sensor chip (GE Healthcare) in HBS, followed by the injection of P0ct. Chip regeneration was performed using a 2:3 (v:v) mixture of 2-propanol and 50 mM NaOH. The used P0ct concentrations were 20 – 2000 nM in HBS, and a single concentration per each lipid capture was studied; all samples were prepared and measured in duplicate. In each run, a single sample was measured twice to rule out instrumental deviation. The binding response as a function of protein concentration was plotted and fitted to the 4-parameter model,

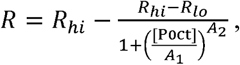

to gain information about association affinity. For kinetic analyses, all association phases (180 s after injection of P0ct) were individually fitted to a one-phase exponential association model using GraphPad Prism 7. The determined *k*_obs_ values were plotted against P0ct concentration and fitted using linear regression to determine *k*_on_ (slope of the curve) and *k*_off_ (Y-intercept of the curve). The values were extracted from two individually fitted datasets: one containing all data (Fitting set 1) as well as one omitting all data points below 350 nM P0ct (Fitting set 2).

### Vesicle turbidimetry and X-ray diffraction

For turbidimetric measurements, SUVs of 0.5 mM DOPC:DOPG (1:1) and DMPC:DMPG (1:1), both with and without supplemented 10% (w/w) cholesterol, were mixed with 0.5 – 10 µM P0ct in duplicate. Light scattering was recorded at 450 nm for 10 min at 25 °C using a Tecan M1000Pro plate reader. The results were analyzed after the observed optical density per time had stabilized.

SAXD experiments were performed to investigate any repetitive structures in turbid samples. 10 and 20 µM P0ct was mixed with SUVs of 1 – 3 mM DMPC:DMPG (1:1) in HBS at ambient temperature and exposed at 25 °C on the EMBL P12 BioSAXS beamline, DESY (Hamburg, Germany). A HBS buffer reference was subtracted from the data. Lipid samples without added P0ct were devoid of Bragg peaks. The peak positions of momentum transfer, *s*, in P0ct-lipid samples were used to calculate mean real-space repeat distances, *d*, in proteolipid structures, using the equation

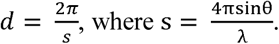

### Atomic force microscopy

Fresh DOPC:DOPS (1:1) SUVs were unrolled on freshly cleaved mica (Ø 1.2 cm) in HBS-Ca (10 mM HEPES, 150 mM NaCl, 2 mM CaCl_2_, pH 7.5), by covering the mica entirely with 0.2 mg ml^−1^ SUVs, followed by a 20-min incubation at 30 °C, and washing twice with HBS-Ca.

The samples were imaged immediately in HBS at ambient temperature using an Asylum Research MFP-3D Bio instrument. TR800PSA cantilevers (Olympus; spring constant (*k*) range 0.59 – 0.68 N m^−1^, resonance frequency 77 kHz) were used in alternative current (AC) mode. Square 256 × 256 pixel scans were acquired from areas between 5 – 20 µm, using a 90° scanning angle and a 0.6 – 0.8 Hz scan speed. The resulting scan images were processed in Igor Pro 6.37.

After confirming the presence of lipid bilayers, 1.8 – 10 µM P0ct was added onto the bilayer samples in HBS. After a 15-min incubation period at ambient temperature, the bilayers were washed twice with HBS, and imaged as above. For each protein concentration, 2 samples were prepared and scanned with identical results. At least 3 different parts per sample were scanned to gain an insight into any sample heterogeneity.

### Electron microscopy

For negative stain EM, 740 µM DMPC:DMPG (1:1) SUVs were mixed with P0ct using protein-to-lipid ratios of 1:25, 1:50, 1:100, 1:200 and 1:500 and incubated at 22 °C for 1 h. EM grids with P0ct or full-length P0 samples were then prepared, stained, and imaged as described ^7, 16^.

For cryo-EM, full-length P0 (0.9 mg/ml) was reconstituted into lipid membranes, after dissolving the lyophilized protein extract in 10 mM sodium phosphate (pH 8.0), 0.4% DM. P0 was mixed with *E. coli* polar lipids or DOPC (both at 5 mg/ml in 2% DM) at lipid-to-protein ratio of 0.5 (w/w) and dialyzed against 10 mM sodium phosphate (pH 8.0) for 5 days at 37 °C. The total ternary mixture volume was 65 µl.

Approximately 3 µl of the P0 reconstituted into *E.coli* polar lipids (∼0.65 mg ml^−1^) were applied onto glow-discharged Quantifoil holey carbon grids (R 1.2/1.3, R 2/2, or R3.5/1, Cu 400 mesh, Quantifoil Micro Tools GmbH, Germany). After 2-s blotting, grids were flash frozen in liquid ethane, using an FEI Vitrobot IV (Vitrobot, Maastricht Instruments) with the environmental chamber set at 90% humidity and a temperature of 20 ºC. Samples were imaged with FEI Titan Krios TEM operated at 300 keV, and images were recorded using a Gatan K2-Summit direct electron detector. Images were collected manually in electron-counting mode at a nominal magnification of ×22,500 and a calibrated pixel size of 1.3 Å. Each image was dose-fractionated to 40 frames (8 s in total, 0.2-s frames, dose rate 6-7 e^-^ / pixel / s). Movie frames were aligned with MotionCorr^56^ and preprocessed on the fly with 2dx_automator^57^. For image processing were used 20 drift-corrected cryo-EM images. Particles were boxed and segmented using e2boxer.py in EMAN2^58^. From all images more than 9000 overlapping, CTF-corrected, segments with size of 160×160 pixels (208 Å × 208 Å) were selected. 2D class averages of P0 between lipid bilayers were calculated with SPRING software^59^.

## Acknowledgements

This work was financially supported by the Academy of Finland (Finland), the Sigrid Jusélius Foundation (Finland), the Emil Aaltonen Foundation (Finland), the University of Oulu Graduate School (Finland), the Norwegian Research Council (SYNKNØYT program), and Western Norway Regional Health Authority (Norway). We thank Dr. Marte Innselset Flydal for practical guidance in using DSC. We gratefully acknowledge the synchrotron radiation facilities and the beamline support at ASTRID2, and EMBL/DESY. We acknowledge the KIT light source for provision of instruments at the beamline UV-CD12 of the Institute of Biological Interfaces (IBG2), and we would like to thank the Institute for Beam Physics and Technology (IBPT) for the operation of the storage ring, the Karlsruhe Research Accelerator (KARA). We also express our gratitude towards the Biocenter Oulu Proteomics and Protein Analysis Core Facility for providing access to mass spectrometric instrumentation and the BiSS facility (Department of Biomedicine, University of Bergen) for the availability of biophysical instrumentation.

## Author contributions

Study design: A.R., S.R., J.K., H.H., P.R., H.S., P.K.

Prepared samples and performed experiments: A.R., S.R., J.K., H.H., A.B., M.M., A.F., R.R., J.B.

Processed and analyzed data: A.R., J.K., S.R., H.H., A.B., P.K.

Original text and figures: A.R., P.K.

Review & editing: A.R., S.R., P.R., J.B., A.S.U., H.S., P.K.

Supervision: P.R., A.S.U., H.S., P.K.

## Data availability

The datasets generated and analyzed during the current study are available from the corresponding author on reasonable request.

